# *In vivo* correction of cystic fibrosis mediated by PNA nanoparticles

**DOI:** 10.1101/2022.01.28.478191

**Authors:** Alexandra S. Piotrowski-Daspit, Christina Barone, Chun-Yu Lin, Yanxiang Deng, Douglas Wu, Thomas C. Binns, Emily Xu, Adele S. Ricciardi, Rachael Putman, Richard Nguyen, Anisha Gupta, Rong Fan, Peter M. Glazer, W. Mark Saltzman, Marie. E. Egan

## Abstract

Cystic fibrosis (CF) is caused by mutations in the cystic fibrosis transmembrane conductance regulator (CFTR) gene. We sought to correct the multiple organ dysfunction of the F508del CF-causing mutation using systemic delivery of peptide nucleic acid gene editing technology mediated by biocompatible polymeric nanoparticles. We confirmed phenotypic and genotypic modification *in vitro* in primary nasal epithelial cells from F508del mice grown at air-liquid interface and *in vivo* in F508del mice following intravenous delivery. *In vivo* treatment resulted in a partial gain of CFTR function in epithelia as measured by *in situ* potential differences and Ussing chamber assays and correction of CFTR in both airway and GI tissues with no off-target effects above background. This is the first report of systemic gene editing for CF. Our data suggest that systemic delivery of PNA NPs designed to correct CF-causing mutations is a viable option to ameliorate the disease in multiple affected organs.

## Introduction

Cystic fibrosis (CF) is an autosomal recessive disorder caused by mutations in the cystic fibrosis transmembrane conductance regulator (CFTR) gene. CFTR encodes a chloride channel key to balancing ion and water secretion and absorption in epithelial tissues. CF patients experience multiorgan dysfunction; for example, CFTR defects cause mucus in several organs to thicken, leading to lung infections, a reduction in lung function, and poor digestive function in the gastrointestinal (GI) tract. While there are over 1700 different CF-causing mutations, the most common mutation is F508del. This is a three base pair deletion in the genomic sequence that results in improper protein folding and impaired transport to the plasma membrane. The recent advancement of modulator therapies designed to mitigate effects of the F508del mutation by increasing transport to the membrane and improving channel function holds great promise, but this regimen requires expensive and continuous treatment. Gene editing approaches, on the other hand, could offer a one-time cure applicable to all CF mutations, including the 10% of patients with rare mutations who are not candidates for modulator therapies.

Efforts to precisely correct genomic mutations that underlie hereditary diseases such as CF for therapeutic benefit have advanced alongside the emergence and improvement of genome editing technologies. These methods fall under two main classes: nuclease-based platforms such as zinc finger nucleases (ZFNs), TALENs, as well as the widely used CRISPR/Cas9 systems including base and prime editors, and oligo/polynucleotide strategies such as triplex-forming oligonucleotides (TFOs).

Programmable RNA-guided Cas9 endonucleases enable efficient genome editing in both cells and organisms, but also cause collateral damage throughout the genome in the form of off-target effects due to nuclease activity, though this can be mitigated by impairing catalytic activity.^1-5^ Moreover, *in vivo* delivery of the large constructs that make up the complex CRISPR/Cas9 system remains challenging.^6^ We have developed a non-nuclease-based approach to gene editing by utilizing endogenous DNA repair stimulated by the binding of peptide nucleic acids (PNAs) to genomic DNA to create a PNA/DNA/PNA triplex structure via both Watson-Crick (WC) and Hoogsteen H-bonding with displacement of the non-bound DNA strand. PNAs have a peptide backbone but undergo base pairing with DNA and RNA.^7^ They also lack intrinsic nuclease activity. Triplex PNA structures can initiate an endogenous DNA repair response mediated by high fidelity nucleotide excision repair (NER) and homology-dependent repair (HR) pathways.^8,9^ When PNAs are introduced with a single-stranded “donor” DNA containing the desired sequence modification, site-specific modification of the genome occurs.^10^ In terms of delivery, PNA and donor DNA are small relative to Cas9 systems and can be readily encapsulated into polymeric vehicles in the form of NPs.^9,10^ We have previously demonstrated the efficacy of PLGA NPs encapsulating PNA and DNA to achieve gene editing both *ex vivo* and *in vivo.* ^11-15^ Here, we demonstrate the utility of PNA NPs for the systemic treatment of F508del CF.

Gene therapies for CF have thus far have had limited success in part due to challenges in delivery to key organs affected by the disease,^16^ namely the lung and gastrointestinal tract. Recent *in vivo* gene editing therapies, primarily employing adenovirus-derived vector systems, focus on targeted correction of the CFTR gene in the airway epithelium.^17-19^ Further, the use of CRISPR/Cas9-based gene editing has been demonstrated for the treatment of CF *in vitro*, with additional studies providing a path for its potential application as a targeted therapy to the lungs *in vivo*.^20-22^ Our own studies have shown that polymeric NPs deliver TFO-based gene editing agents to the lung, with phenotypic correction.^14^ However, CF is a systemic disease with multiple affected organs that could potentially benefit from gene correction therapies; while the *in vivo* approaches for remedying CF via gene therapy have shown promise, most recent studies remain limited to a local scope and do not tackle the systemic breadth of the disease.^21^ Here we utilize PNA NPs for systemic delivery to correct the F508del mutation. We demonstrate their use both *in vitro* in primary cells grown in a physiologically relevant air-liquid interface (ALI) culture model and *in vivo* in mice homozygous for the F508del mutation. This is the first report of systemic therapeutic gene editing to correct CF-causing mutations *in vivo*.

## Results

To determine the feasibility of systemic gene editing as a therapeutic approach for CF-treatment, we utilized a murine CF model homozygous for the F508del mutation in Exon 11. Based on our previous work with local intranasal delivery of PNA NPs in these mice,^14^ we designed tail-clamp PNA molecules that bind near the mutation site with homopurine/homopyrimidine stretches (**Figure 1a**). Here, we incorporated modified PNA monomers with a mini-polyethylene glycol group at the γ-position into the PNA sequence in the Hoogsteen domain. γPNAs exhibit enhanced DNA binding;^23^ we have previously demonstrated corresponding elevations in gene correction *ex vivo* and *in vivo* using these γ-modified PNAs in a β-thalassemia mouse,^13^ and also in human CF bronchial epithelial cells (CFBEs) homozygous for the F508del mutation.^24^ We also designed a single-stranded correcting donor DNA template to introduce 3 nucleotides consistent with the wild-type genomic sequence. Both PNA and donor DNA molecules were loaded into polymeric NPs in a 2:1 molar ratio (**Figure 1b**) consisting of poly(lactic-co-glycolic acid) (PLGA) formulated using a double emulsion solvent evaporation technique as described previously.^9,13^ The resulting NPs were spherical and ~250 nm in diameter, as characterized by dynamic light scattering (DLS) (**Supplementary Table 1**) and scanning electron microscopy (SEM) (**Supplementary Figure 1**). These NPs were administered both *in vitro* in ALI cultures of primary airway epithelial cells and *in vivo* via systemic intravenous (IV) injection to F508del-CFTR mice.

**Figure 1.**
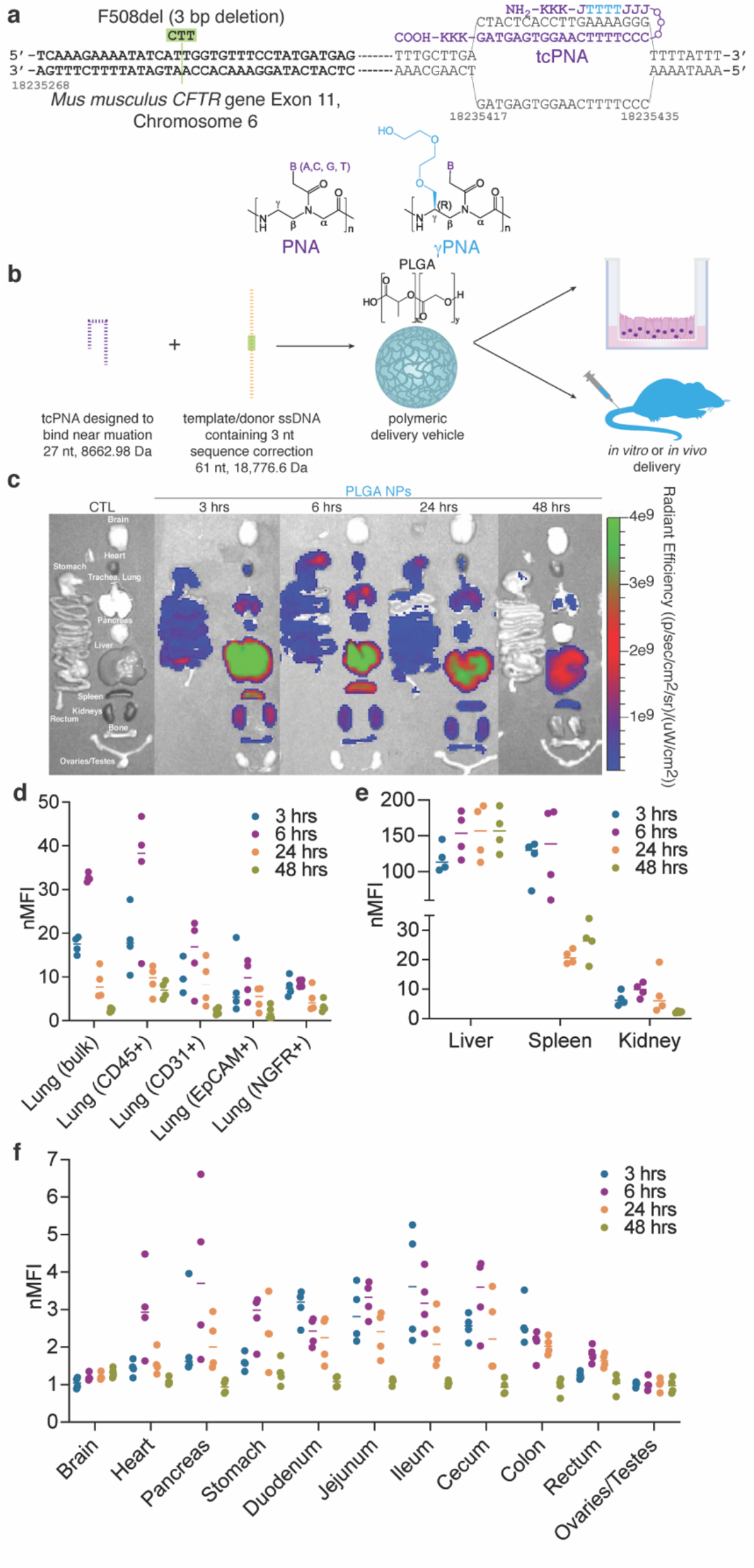
PNA-based gene editing agents can be encapsulated into PLGA NPs, which exhibit accumulation in lung and GI tract following systemic IV administration. (a) Schematic of PNA design to correct F508del CFTR indicating the incorporation of γPNA monomers and the formation of the PNA/DNA/PNA triplex. (b) PNA and donor DNA *in vitro* and *in vivo* delivery strategy: encapsulation into polymeric PLGA NPs. (c) Representative IVIS images indicating biodistribution of Cy5-conjugated PLGA NPs at 3, 6, 24, and 48 hours post-IV administration *in vivo* compared to an untreated control animal (CTL). (d) Flow cytometry mean fluorescence intensity values normalized to untreated control animals (nMFI) for homogenized bulk lung and specific cell types (CD45+ macrophages, CD31+ endothelial cells, EpCAM+ epithelial cells, and NGFR+ basal cells) at 3, 6, 24, and 48 hours post-IV administration of Cy5-conjugated PLGA NPs *in vivo*. (e) Flow cytometry mean fluorescence intensity values normalized to untreated control animals (nMFI) for homogenized bulk liver, spleen and kidney at 3, 6, 24, and 48 hours post-IV administration of Cy5-conjugated PLGA NPs *in vivo*. (f) Flow cytometry mean fluorescence intensity values normalized to untreated control animals (nMFI) for homogenized bulk brain, heart, pancreas, stomach, duodenum, jejunum, ileum, cecum, colon, rectum, and ovaries/testes at 3, 6, 24, and 48 hours post-IV administration of Cy5-conjugated PLGA NPs *in vivo*.

As CF is a systemic disease with a heavy burden on both airway and GI epithelia, we first tested whether PLGA NPs could reach these organs when administered systemically. In prior work, we have found that NP size in particular is important for NP accumulation in multiple tissue types *in vivo* following intravenous administration.^25^ We assessed the biodistribution of Cy5-conjugated PLGA NPs at several timepoints after systemic IV administration. Whole organ fluorescence was captured using an In Vivo Imaging System (IVIS) and indicated high levels of NP accumulation in the lung at 3, 6, and 24 hrs post-administration (**Figure 1c**). Similar to previous reports of PLGA biodistribution, we also observed high accumulation in the liver and spleen,^26^ as well as lower levels of accumulation other organ systems of interest for CF, including the GI tract. It is important to note, however, that IVIS analysis cannot determine which cell types have taken up the NPs, and so we also performed flow cytometry analyses. Consistent with the IVIS data, we observed an increase in lung NP accumulation by flow cytometry at 3 and 6 hrs post-administration (**Figure 1d**). Notably, we observed NP uptake in multiple cell types in the lung, including CD45+ macrophages, CD31+ endothelial cells, EpCAM+ epithelial cells, and NGFR+ basal cells. Flow cytometry analyses of other tissues including liver, spleen, kidney, brain, heart, pancreas, stomach, duodenum, jejunum, ileum, cecum, colon, rectum, and ovaries/testes were also consistent with the IVIS data. Lastly, we confirmed uptake in key tissues relevant to *in vivo* CFTR function assessment, namely the nasal epithelium and rectum, by microscopy (**Supplementary Figure 2**). In summary, these data suggest that PLGA-based NPs are able to reach target organs and cell types relevant to CF treatment after IV delivery. While the precise mechanisms of intracellular trafficking of PLGA NPs remain unclear, we and others have shown that uptake occurs primarily by adsorptive-type endocytosis.^27,28^ Once inside cells, NPs are rapidly shuttled into the endolysosomal pathway and escape from the late endosome into the cytoplasm. In prior work with PNA/DNA NPs, we have shown that PLGA NPs readily associate with and are taken up by various cell types, and that they accumulate in the perinuclear region.^29^

We next sought to develop a physiologically relevant *in vitro* model in which to test the therapeutic activity of our PNA NPs, taking advantage of ALI systems commonly used in CF research. In this case, we cultured primary nasal epithelial cells (NECs) to more closely mimic the *in vivo* environment. There are several benefits of this experimental system: 1) these cells can be expanded using feeder cells (fibroblasts) and provide an *in vitro* model in which we can test our gene editing reagents targeting the murine CFTR genomic locus; 2) these are primary cells which mature into pseudostratified epithelia with multiple cell types; 3) to culture these NECs, we adapted clinically relevant methods described to culture human nasal epithelial samples, ^30^ and 4) this system and protocol are comparable to those used for human nasal epithelial cell culture. To develop the culture system, primary NECs were isolated from F508del mice (**Figure 2a**) and expanded using protocols recently described for the expansion of primary human airway epithelial cells.^30^ After expansion, cells were seeded on permeable supports in transwell inserts and transitioned to ALI over the course of several weeks. These cells formed pseudostratified epithelial layers similar to the epithelial structure *in vivo* (**Figure 2b**), and were primarily composed of basal cells when seeded (**Figure 2c**), with multiple cell types present in the mature cultures. To further characterize the model system, we performed single-cell RNA sequencing analysis (scRNAseq) on ~10^5^ primary nasal epithelial cells cultured at ALI using cell-type markers (ex. transcription factors and surface molecules) described in recent reports.^31,32^ The scRNAseq data was visualized using a graph-based algorithm (Uniform Manifold Approximation and Projection, UMAP)^33,34^ to facilitate the identification of distinct cell types. Cells were partitioned into distinct populations with unique gene expression signatures (**Figure 2d**). Several cell types were identified, reminiscent of the *in vivo* environment: basal cells (the presumptive stem cell in the airway epithelium)^35^, cycling basal cells, ciliated cells, and club cells.

**Figure 2.**
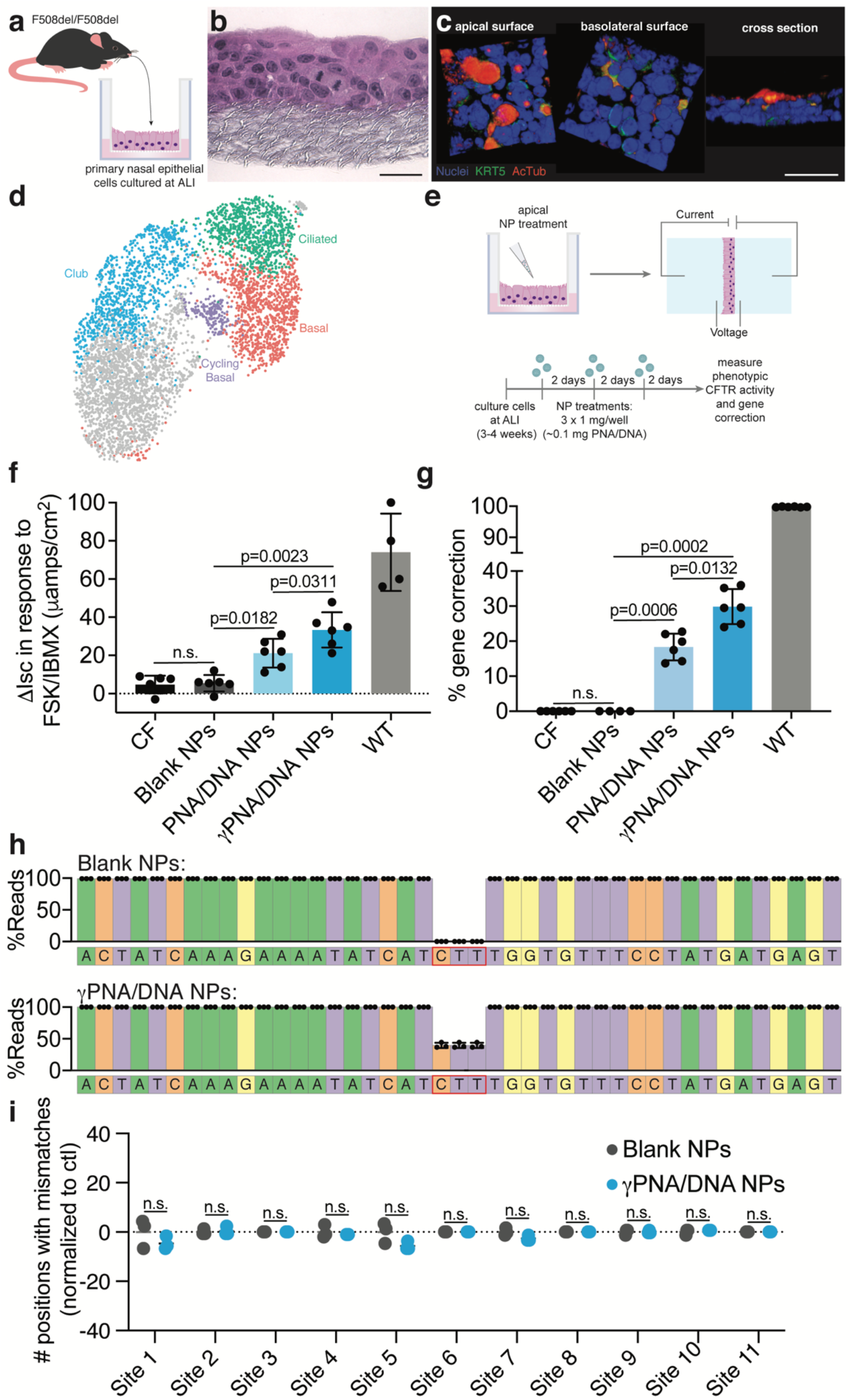
PNA NP treatment results in phenotypic and genotypic CFTR correction in physiologically relevant primary NEC ALI cultures consisting of multiple cell types. (a) Schematic of primary NEC isolation. (b) H&E stained parrafin sections of NEC ALI cultures. Scale bar, 50 μm. (c) 3D renderings of confocal images of NEC cultures indicating a pseudostratified epithelium containing basal cells (Krt5) and ciliated cells (AcTub, acetylated α-tubulin). Scale bar, 50 μm. (d) UMAP plot of scRNAseq data for primary NEC ALI cultures indicating distinct cell types (basal, cycling basal, ciliated, and club) (n= 3 cultures). (e) Schematic of NP treatment scheme in primary NEC ALI cultures. Cells were treated with 3 doses of 1 mg Blank or PNA/DNA PLGA NPs. PNA/DNA NPs contain ~2 μg/0.2 nmol of PNA and ~2 μg/0.1 nmol DNA per mg. (f) Phenotypic Δ*I_sc_* measurements following treatment of primary NEC ALI cultures with Blank, PNA/DNA, or γPNA/DNA NPs compared to CF or wild-type (WT) controls. (g) ddPCR gene correction assessments of NP-treated primary NEC ALI cultures. (h) Deep sequencing read analysis around the target site of NP-treated primary NEC ALI cultures. (i) Deep sequencing off-target analysis of Blank or γPNA/DNA NP-treated NEC ALI cultures at 11 genomic sites with partial PNA binding site homology displayed as the number of positions with mismatches to the reference sequence normalized to untreated control samples.

We used this primary NEC ALI culture model to test the hypothesis that γ-modified PNAs could mediate enhanced gene correction compared to unmodified PNAs. PNA and γPNA NPs were administered to the primary NEC ALI cultures and assessed for both phenotypic and genotypic CFTR correction. To mimic the multi-dose *in vivo* treatment schemes that we have used in previous studies, each culture received a dose of 1 mg NPs every 2 d for a total of 3 doses (**Figure 2e**). Two days after the last treatment, CFTR activity was assessed via Ussing chamber measurements. We monitored CFTR-mediated ion transport as the change in short-circuit current (Δ*I_sc_*) in response to a CFTR-stimulating cocktail of forskolin and IBMX. Representative traces are shown in **Supplementary Figure 3**. Compared to untreated and blank NP-treated samples, we observed significant increases in Δ*I_sc_* for both PNA and γPNA NP treated samples (**Figure 2f**). The Δ*I_sc_* recorded for γPNA NP-treated samples was significantly higher than the Δ*I_sc_* for PNA NP-treated samples. After Ussing measurements, we next assessed the extent of genotypic correction giving rise to these phenotypic changes. To reduce potential PCR bias, we developed a droplet digital PCR (ddPCR) assay to quantify the percentage of modified CFTR alleles in the NEC cultures (**Supplementary Figure 4**; **Figure 2g**). Consistent with the phenotypic changes we measured in the Ussing chamber, in apically treated NEC cultures, we observed ~18% correction after PNA NP treatment and ~30% correction after γPNA NP treatment. We also performed next generation sequencing (NGS) and did not observe unintended indels at the target site (**Figure 2h**). Further, in 11 genomic sites with partial homology to the PNA or donor DNA binding sites identified using BLAST (**Supplementary Table 2**), the off-target mutation/error rates were similar to blank NP-treated controls with minimal variation from the reference sequence (**Figure 2i**). Together, these results suggest that PNA-based gene editors can produce both phenotypic and genotypic correction. Further, γPNA NPs are significantly more effective for gene correction compared to PNA NPs in primary airway epithelial cells with a large basal cell population. These measurements are consistent with results we have observed in human CFBE cells and with inhalational delivery of PNA vs. γPNA NPs in the F508del mouse.^24^

We next tested for phenotypic and genotypic correction of the F508del CF mutation in a murine disease model homozygous for the 3-bp deletion (F508del-CFTR). PLGA NPs encapsulating γPNA and donor DNA designed to correct the F508del mutation were formulated and F508del-CFTR mice were treated with a 2 mg NP resuspension administered IV for four treatments over the course of 2 wk (**Figure 3a**). Both blank and γPNA/DNA NPs are well tolerated without indications of toxicity or inflammation (**Supplementary Table 3, Supplementary Figure 5,6**). Two weeks after the last treatment, *in vivo* phenotypic correction of the F508del mutation was assessed using non-invasive assays to detect CFTR activity *in vivo*: nasal potential difference (NPD) and rectal potential difference (RPD). The PD assay is a useful method to study ion transport if serial measurements are required.^36^ Nasal and rectal epithelia in wild-type mice exhibit a robust cyclic AMP (cAMP)-stimulated chloride efflux, whereas CFTR dysfunction results in a lack of activation of cyclic AMP-stimulated chloride flux. After one dosing round of IV-delivered γPNA NPs, we observed a partial amelioration of the impaired response to cAMP stimulation in the nasal epithelia, with some mice exhibiting hyperpolarized (more negative) ΔNPD in response to CFTR stimulation into the wildtype range (**Figure 3b**), though the degree of hyperpolarization was variable with a subset of mice not responding to treatment. Untreated control and blank NP-treated mice exhibited typical CF NPD responses: a lack of hyperpolarization or a depolarization in response to a cAMP stimulation cocktail which is consistent with the absence of CFTR activity. Representative raw NPD traces are shown in **Supplementary Figure 7**. Topical intranasal delivery of the same γPNA and donor DNA reagents using modified PLGA/PBAE/MPG NP formulations (as we have described previously) resulted in NPD responses closer to and not significantly different from wild-type values, consistent with our previous observations (**Supplementary Figure 8**).^14^ The RPD responses to cAMP stimulating agents of systemically γPNA NP-treated mice were not significantly different from untreated and blank NP-treated controls, though a few mice exhibited hyperpolarization into the wild-type range (**Figure 3c**). An examination of pre-treatment and post-treatment NPD responses to cAMP stimulation for each mouse reveals an average percent change of −118% in the ΔNPD for γPNA NP-treated mice, whereas the average percent change in ΔNPD for blank NP-treated mice was +28% compared to baseline measurements (**Figure 3d,e**). A similar analysis of RPD responses shows an average percent change of −50% in ΔRPD for γPNA NP-treated mice and an average percent change of −6% in ΔRPD for blank NP-treated mice (**Figure 3f,g**). To assess the longevity of treatment responses, we performed additional NPD and RPD measurements for a subset of Blank NP and γPNA NP-treated mice two weeks after the first post-treatment measurements. We observed an attenuation in the NPD response in the second measurement for the two mice that responded to γPNA NP-treatment but no change in response for one mouse that did not exhibit a change in NPD following the first round of treatment (**Figure 3h**). This attenuation in response is consistent with reports by others studying the longevity of gene editing, and specifically HDR repair events, with CRISPR/Cas9.^37^ Following the second post-treatment measurements, we re-treated each mouse with another round of four NP doses and measured the NPD response two weeks after the last treatment. After the second treatment round, the mice that had shown a response after the first treatment round exhibited NPD hyperpolarization in response to cAMP stimulation cocktail again, whereas the mice that did not respond to the first round of treatment did not respond to the second treatment. We observed similar trends for RPD measurements performed on the same mice (**Figure 3i**). Blank NP-treated animals exhibited similar NPD measurements throughout the study, consistent with measurements of untreated animals from a variety of CF mouse models.^36^

**Figure 3.**
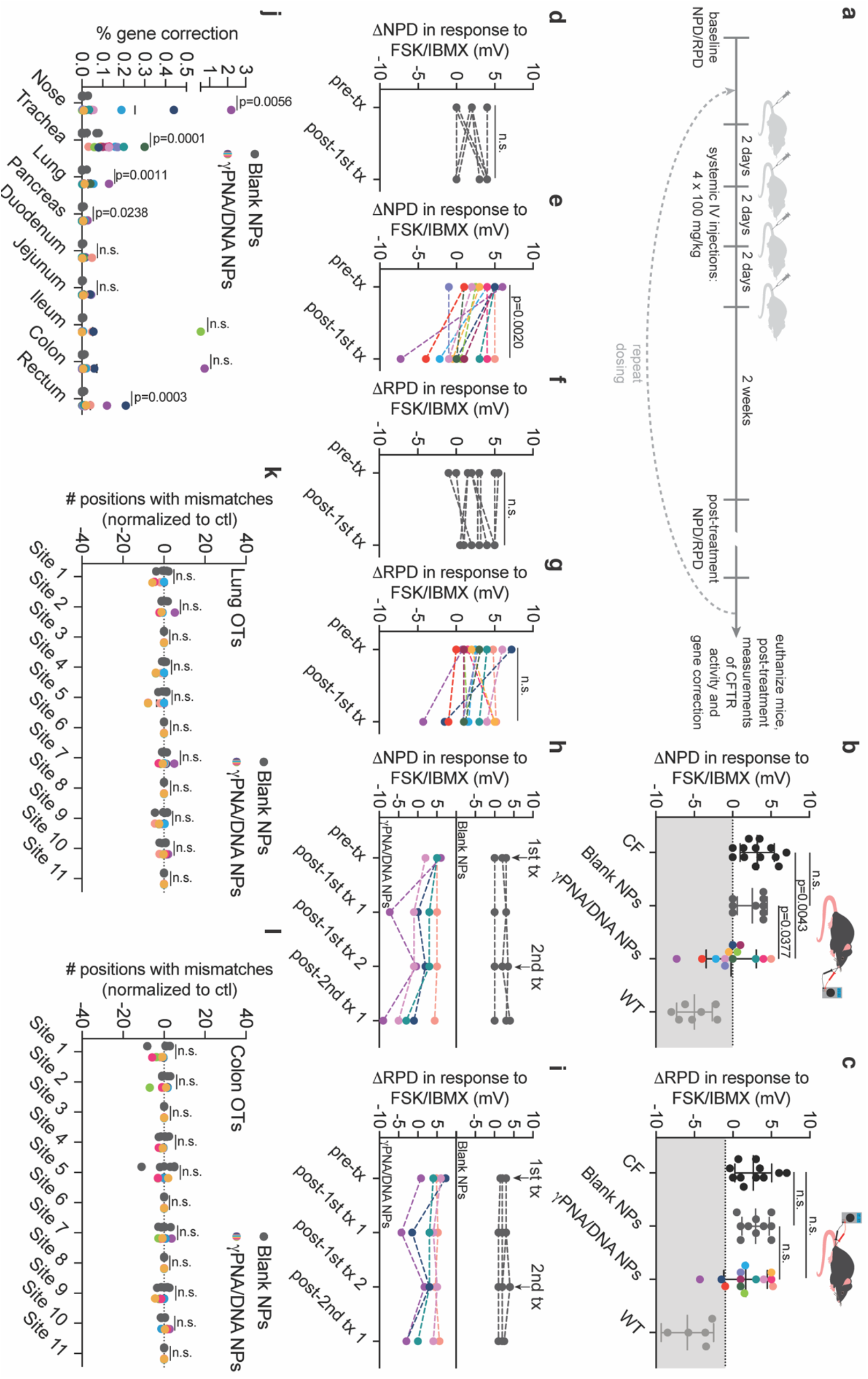
Functional and genotypic correction of CFTR *in vivo* following PNA NP administration. (a) Schematic of *in vivo* NP dosing scheme. (b) Nasal potential difference (NPD) measurements following either 4 x 2 mg blank NP (dark grey circles) or 4 x 2 mg γPNA/DNA NP (multicolored circles) treatment with CF (black circles) and wildtype (light grey circles) controls. γPNA/DNA NPs contain ~2 μg/0.2 nmol of PNA and ~2 μg/0.1 nmol DNA per mg; each animal received ~0.2 mg/kg of PNA and donor DNA per dose. Each color represents a different mouse in the γPNA/DNA NP-treated cohort. Grey region indicates wild-type range. (c) Rectal potential difference (RPD) measurements following either Blank NP (dark grey circles) or γPNA/DNA NP (multicolored circles) treatment with CF (black circles) and wildtype (light grey circles) controls. Each color represents a different mouse in the γPNA/DNA NP-treated cohort. Grey region indicates wild-type range. (d) Pre- and post-treatment NPD measurements for F508del-CFTR mice treated with Blank NPs. (e) Pre- and post-treatment NPD measurements for F508del-CFTR mice treated with γPNA/DNA NPs. Each color represents a different mouse. (f) Pre- and post-treatment RPD measurements for F508del-CFTR mice treated with Blank NPs. (g) Pre- and post-treatment RPD measurements for F508del-CFTR mice treated with γPNA/DNA NPs. Each color represents a different mouse. (h) Serial NPD measurements performed over the course of two treatment rounds for a subset of animals in the γPNA/DNA NP-treated and Blank NP-treated cohorts. Arrows indicate treatment round initiation. (i) Serial RPD measurements performed over the course of two treatment rounds for a subset of animals in the γPNA/DNA NP-treated and Blank NP-treated cohorts. Arrows indicate treatment round initiation. (j) Gene correction levels measured by ddPCR at the F508del locus for airway and GI organs from mice treated with either blank NPs or γPNA/DNA NPs. Deep sequencing off-target analysis of Blank or γPNA/DNA NP-treated F508del mice (k) lungs and (l) colons at 11 genomic sites with partial PNA binding site homology displayed as the number of positions with mismatches to the reference sequence normalized to untreated control samples.

In addition to phenotypic measurements of CFTR correction *in vivo*, we also assessed gene correction in bulk tissues by ddPCR in several organs. We observed editing in tissues that make up the airways and GI tract, including the nasal epithelium (up to ~2%), trachea (up to ~0.3%), lung (up to ~0.1%), ileum (up to ~0.5%), colon (up to ~0.7%), and rectum (up to ~0.2%) (**Figure 3j**). There was a high degree of heterogeneity and variability in gene correction measurements amongst the treated mice, consistent with CRISPR-based editing approaches.^38^ However, the degree of editing for each mouse was consistent with the degree of electrophysiological response in NPD and RPD assays. NGS was also used to detect off-target editing events in the lung and colon (**Figure 3k,l**) with a focus on genomic regions with partial homology to the PNA or donor DNA binding sites. Detected mismatches were equivalent to those observed for control blank NP-treated samples with low variation from the reference sequence.

We further assessed phenotypic disease amelioration by measuring cell counts in the bronchoalveolar lavage (BAL) fluid of treated mice, which typically includes alveolar macrophages and neutrophils (**Figure 4a**).^39^ We observed consistently lower cell counts in γPNA/DNA NP-treated mice, suggesting partial restoration of CFTR function. In addition, we assessed CFTR function in treated mice by performing Ussing chamber measurements of Δ*I_sc_* in response to CFTR stimulation as described above on epithelial tissues *ex vivo*. Tissues (nasal epithelium, rectum, distal colon, ileum, duodenum, and jejunum) were immediately dissected after euthanasia and mounted into Ussing chambers (**Figure 4b**). We recorded Δ*I_sc_* values for tissues for intact epithelia where possible. For several tissues including nasal epithelium, rectum, and distal colon, γPNA/DNA NP-treated mice exhibited Δ*I_sc_* measurements significantly different from both blank NP-treated animals and CF controls (**Figure 4c-h**). Further, we observed again that the mice with more robust *in vivo* phenotypic responses had larger Δ*I_sc_* values.

**Figure 4.**
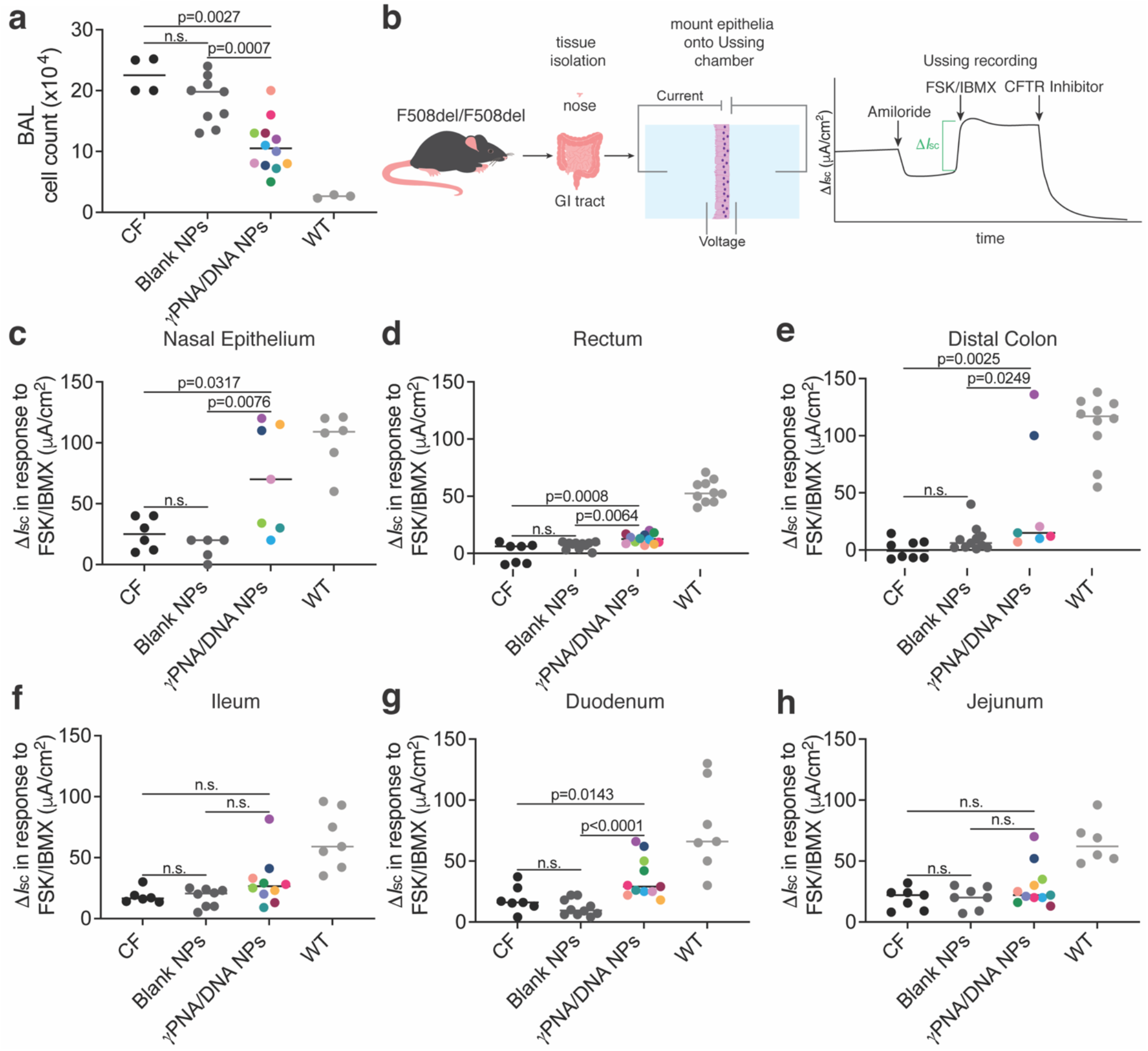
Functional correction of CFTR *ex vivo* following PNA NP administration. (a) BAL cell counts following either blank NP (dark grey circles) or γPNA/DNA NP (multicolored circles) treatment with CF (black circles) and wildtype (light grey circles) controls. γPNA/DNA NPs contain ~2 μg/0.2 nmol of PNA and ~2 μg/0.1 nmol DNA per mg; each animal received ~0.2 mg/kg of PNA and donor DNA per dose. Each color represents a different mouse in the γPNA/DNA NP-treated cohort. (b) Schematic of *ex vivo* tissue isolation and Ussing chamber assay. Phenotypic Δ*I_sc_* measurements in (c) nasal epithelium, (d) rectum, (e) distal colon, (f) ileum, (g) duodenum, and (h) jejunum following either 4 x 2 mg blank NP (dark grey circles) or 4 x 2 mg γPNA/DNA NP (multicolored circles) treatment with CF (black circles) and wildtype (light grey circles) controls. Each color represents a different mouse in the γPNA/DNA NP-treated cohort.

## Discussion

The therapeutic approach for CF described here combines non-nuclease-based PNA gene editing technology with systemic delivery of biocompatible polymeric NPs to achieve gene correction in multiple tissue types *in vitro* and *in vivo*. Improved PNA molecules containing structure-modifying γ-substitutions demonstrated phenotypic correction and gene editing without unintended indel formation when co-delivered with donor DNA *in vitro*, to primary nasal epithelial cells isolated from mice harboring the F508del-associated CF mutation, and *in vivo* in the same mouse model. Enhanced gene editing with modified PNAs is consistent with our prior work in another disease context.^13^ We believe this is the first evidence of *in vivo* systemic delivery of gene editing agents to correct a CF-causing mutation in multiple organs.

From a safety perspective, both the therapeutic PNA/donor DNA combination as well as the polymeric delivery vehicle did not exhibit detectable adverse effects. Low levels of off-target genomic mutations and well-tolerated systemic delivery are paramount to clinical translation of gene editing therapeutics. The intended correcting 3-bp CTT insertion was detected at the target site by both ddPCR and deep sequencing, with no unintended outcomes in the form of indels created in the flanking sequences. While much progress has been made with regard to methods for determining genome-wide off-target mutations for nuclease-based editing technologies,^40-43^ our ability to assess the potential off-target effects of PNA-based editing technology remains limited to sequence homology. As expected for PNA molecules with sequence-specificity, deep sequencing of multiple sites with partial homology to the PNA binding site revealed no off-target effects above background mutation/read error rates *in vitro* in treated nasal epithelial cells and *in vivo* in the lungs and colons of treated mice. Additional concerns with systemically delivered gene editing therapeutics are the potential adverse consequences of the editors or the delivery vehicles.^44^ PNA NPs did not exhibit systemic toxicity as evidenced by serum chemistry analyses compared to untreated or blank NP controls. Lastly, particularly for intravenously injected therapeutics, the risk of on- or off-target or activity in inappropriate tissues highlights the need to ensure proper tissue tropism. While we observed widespread NP biodistribution to multiple tissues apart from airway and GI epithelia, we did not observe any pathology in tissues with particularly high NP accumulation, including the liver and spleen.

The work presented here provides a foundation for systemic *in vivo* gene editing to correct CF with PNA NPs. There are few studies to date that have described the systemic delivery of therapeutic gene editing agents, especially in disease animal models. One recent study reported 5-7.3% indels produced by Cas9 RNPs encapsulated in lipid nanoparticles in the liver following intravenous delivery in wild-type mice to edit the PCSK9 locus.^45^ While the response to PNA/DNA NP treatment across the cohort of mice studied was variable and the frequency of edited alleles was low, we observed both phenotypic and genotypic correction of extrahepatic tissues *in vivo*. Importantly, elevated gene correction levels correlated well with performance in phenotypic assays. It has been estimated in several studies that correcting CFTR in ~5-15% of cells in epithelia should restore transepithelial chloride secretion to near wild-type levels, suggesting that *in vivo* correction of even a fraction of cells could provide therapeutic benefit.^20,46-50^ While we did not observe editing in that range in our systemic *in vivo* studies, our results do suggest that even modest levels of editing can result in partial restoration of chloride transport. Compared to systemic delivery, local intranasal delivery of γPNA/DNA NPs resulted in phenotypes more consistently in the wild-type range. In light of these findings, local delivery to the airway epithelia might be combined with systemic delivery to maximize phenotypic correction across multiple organs affected by CF. Systemic delivery in particular can be further improved as a therapeutic strategy. For example, enhancements in vehicle design could be used to increase efficacy, including surface modifiers to target specific cell types as well as the use of different polymeric materials and formulation techniques to vary nanoparticle size and composition with the ultimate goal of modulating encapsulation efficiency and tissue tropism, particularly to the GI tract. Beyond nanoparticle engineering, a more in-depth parameter exploration of dosing and frequency of administration would also elucidate the limits of this approach.

The longevity of treatment response is also an important consideration for translation of gene editing therapeutics. We observed an attenuation in treatment response over time that was reversed upon additional treatment rounds. Optimization of the delivery to target stem cells will be key to obtaining a one-time cure.

## Methods

### Materials

Boc-protected PNA monomers were purchased from ASM Research Chemicals. Mini-PEG γPNA monomers were prepared from Boc-2(2-(2-methoxyethoxy)ethyl)-L-serine as described previously.^13,23^ PNA oligomers were synthesized on solid support (MBHA (4-methylbenzhydrylamine) resin) using Boc chemistry^23^ and purified using HPLC. The tcPNA sequences used in this study was: H-KKK-J<u>TTTT</u>JJJ-OOO-CCCTTTTCAAGGTGAGTAG-KKK–NH^2^, with the underlined portion indicating the positions of γPNA monomers for the γPNA version of the sequence; K, lysine; J, pseudoisocytosine (a cytosine analog for improved PNA/DNA/PNA triplex formation at physiologic pH); O, 8-amino-2,6,10-trioxaoctanoic acid linkers connecting the Hoogsteen and Watson–Crick domains of the tcPNAs. The Donor DNA oligonucleotide (61 nt) was synthesized by Midland Certified Reagent (Midland, TX) and end-protected with three phosphorothioate internucleotide linkages at both the 5’ and 3’ ends and purified by reverse phase HPLC. This antisense donor DNA sequence matches the corresponding region in mouse CFTR exon 11 with the correcting 3 nt insertion underlined:5’T(s)C(s)T(s)TATATCTGTACTCATCATAGGAAACACCA<u>AAG</u>ATAATG TTCTCCTTGATAGTACC(s)C(s)G(s)G3’. PLGA polymer (50:50 DL-PLG, inherent viscosity 0.55-0.75 dL/g) was purchased from Lactel and used as received. Dichloromethane (DCM, HPLC grade, 99+%) was purchased from Sigma Aldrich. DiD ((DiIC18(5); 1,1′-dioctadecyl-3,3,3′,3′-tetramethylindodicarbocyanine, 4-chlorobenzenesulfonate salt) dye was purchased from Biotium and dissolved in DMSO at 10 mg/mL prior to use. Heparin (1000 USP/mL) was purchased from Cardinal Health. Isoflurane was purchased from Sigma Aldrich. Fisherbrand Superfrost Microscope slides were purchased from ThermoFisher Scientific.

### NP formulation and characterization

PLGA NPs encapsulating DiD were formulated using a single oil-in-water (o/w) emulsion solvent evaporation technique as previously described.^51^ Briefly, 50 mg of polymer was dissolved in 900 μL of DCM overnight. 100 μL of DiD dye at 2.5 mg/mL in DMSO was added to the dissolved polymer immediately prior to formulation (0.5% (w/w)). The polymer and dye solution was added dropwise under vortex into 2 mL of 5% (w/v) low MW polyvinyl alcohol (PVA) solution and sonicated with a probe tip sonicator to form an oil-in-water single emulsion and then diluted into 10 mL of 0.3% (w/v) PVA solution while mixing. The remaining organic solvent was evaporated using a rotary evaporator. The NPs were then washed twice in nuclease-free diH_2_O by centrifugation at 16,000 × g to remove excess PVA. NP size and zeta potential were measured via dynamic light scattering (DLS). NP morphology was visualized by scanning electron microscopy (SEM). Blank PLGA NPs and NPs loaded with PNA and donor DNA were formulated using a water-in-oil-in-water (w/o/w) double-emulsion, solvent evaporation technique as described previously.^13^ PNA and DNA were encapsulated at a 2:1 ratio with 2 nmol of PNA and 1 nmol of DNA loaded per mg of polymer. Blank NPs were loaded with 150 μL of nuclease-free water, and PNA/DNA NPs were loaded with 100 μL of PNA and 50 μL of donor DNA each at 1 mM in nuclease-free diH_2_O. NP formulations were resuspended in nuclease-free diH_2_O with 30 mg trehalose per 50 mg initial polymer weight, flash frozen at −80°C, lyophilized for at least 48 hours, and stored at −20 °C.

### Isolation of primary nasal epithelial cells

Primary nasal epithelial cells from mice homozygous for the F508del mutation (fully backcrossed C57/BL6 background) were isolated based on similar protocols described for human cells.^30,52-54^ Briefly, noses were dissected and the nasal epithelium was digested with a protease mixture (1% protease, 0.01% DNase) for 2 hours with agitation at 4 °C. Nasal epithelial cells were then collected and filtered through a 70 μm cell strainer and washed twice by centrifugation (1500 rpm, 10 minutes). Cells were collected by centrifugation and resuspended in F media as previously described^30^ containing 5 μM Y-27632 and cultured on type I collagen (Purecol; Advanced BioMatrix) coated tissue culture dishes with irradiated feeder cells. Differential trypsinization was used to separate feeder and epithelial cells during passaging as previously described.^53^

### ALI cell culture, NP treatments, and Ussing measurements

To initiate air-liquid interface (ALI) cultures, ~1.5 x 10^6^ cells were seeded on permeable polycarbonate supports with 0.4 μm pore size (Corning Costar 3801) in 250 μL of cell culture medium. These 6-well plate inserts had a diameter of 12 mm and were coated with type I collagen (RatCol; Advanced BioMatrix) and exposed to UV radiation overnight. 3 mL of culture medium was added to each well below the inserts. Cells were fed both apically and basolaterally three times per week over the course of 4 weeks after which they were transitioned to ALI and fed only from the bottom. For *in vitro* PNA/DNA NP experiments, cells at ALI were treated either apically or basolaterally every two days with 1 mg of NPs (containing ~2 μg/0.2 nmol of PNA and ~2 μg/0.1 nmol DNA) resuspended in 250 μL of cell culture medium by water bath sonication and vortexing for a total of three treatments. Ussing measurements were performed two days after the last treatment. Ussing experiments with ALI cultures were performed as previously described using an Easy Mount Ussing Chamber System (Physiologic Instruments).^55^ Briefly, chambers were heated to 37 °C and current and voltage electrode tips were prepared by partially filling with 3% agar in 3M KCl and then backfilling with 3M KCL solution. The transwell inserts containing ALI cultures were loaded into P2300 snapwell chambers and filled with 6 mL of Kreb’s Bicarbonate Ringers Solution at 37 °C (140 mM Na^+^, 120 mM Cl_2_, 5.2 mM K^+^, 1.2 mM Ca^2+^, 1.2 mM Mg^2+^, 25 mM HCO_3_^2−^, 2.4 mM HPO_4_^2−^, 0.4 mM H_2_PO_4_^2−^, and 10 mM glucose at pH 7.4) and allowed to equilibrate for 20 minutes. A mixture of 95% O_2_ and 5% CO_2_ gas was bubbled through the solutions. Current-clamped Ussing experiments were performed with a bidirectional pulse of 1 μA of current for 3 seconds every 60 seconds, with transepithelial voltages and membrane resistances measured during each pulse. Amiloride (100 μM) was added apically with a 10-minute equilibration and subsequently maintained. Short-circuit current (*I_sc_*) was then calculated using Ohm’s Law when the apical chamber solution was replaced with a 0 Cl^−^ solution containing forswkolin (10 μM) and IBMX (1 mM) with a 15-minute equilibration. CFTR-specific inhibitor 172 (20 μM) was added apically after 20 minutes, followed by bilateral replacement of solution with Kreb’s Bicarbonate Ringers solution.

### Single-cell RNA-seq (scRNA-seq) analysis

The scRNA-seq was performed using scFTD-seq.^56^ Briefly, a polydimethylsiloxane (PDMS) microwell array chip was used for co-isolating single cells and uniquely barcoded mRNA capture beads. The design of the microwell arrays has been described previously.^56^ Each microchip has up to 25,000 wells and allows for barcoding of ~2,500 cells in a single run to prepare a sequencing library. Library preparation and sequencing steps follow the same protocols as outlined previously.^56^ Transcriptome alignment to a reference transcriptome of the corresponding species including barcode/UMI identification and collapsing was performed as described in the scFTD-seq methods.^56^ The data analysis was performed using Seurat v3.1 on the normalized and log-transformed gene expression data.^57,58^ A total of 5,896 individual cells were used for single-cell data analysis. Cells with low expression of genes (<100 genes), high expression of genes (>2,000), and a high percentage of mitochondrial genes (>15%) were digitally filtered out, resulting in 5,302 single cells used for the subsequent analysis. After identifying the top 2,000 variable genes, we performed principal component analysis (PCA) as well as unsupervised clustering using Uniform Manifold Approximation and Projection (UMAP),^33^ which was implemented in Seurat.^57,58^ For cell type identification, support vector machine (SVM) with linear kernel was used.^59^ In brief, the SVM model was first trained with labeled single cell data from mouse tracheal epithelial cells^31^ with 98.6% accuracy. Then, the trained model was applied to predict the cell types in this work.

### Immunofluorescence and microscopy

To prepare parrafin-embedded NEC sections, cells were immediately fixed in 10% normal buffered formalin for 24 hours at room temperature before being transferred to PBS and kept at 4 °C until paraffin embedding. For immunofluorescence, 5 μm sections were baked at 60 °C for 1 hour and deparrafinized using standard procedures. Antigen retrieval was performed by steaming with citrate buffer (pH 6; Abcam), after which sections were rinsed with DPBS and blocked with 10% normal goat serum (NGS) for 30 minutes at room temperature. Primary antibodies in blocking buffer were added overnight at 4 °C. Sections were then washed three times with DPBS for 5 minutes and secondary antibody diluted in blocking buffer was added for one hour at room temperature. Sections were washed again with PBS three times for 5 minutes. Hoechst 33342 diluted 1:1000 in PBS was then added for 30 seconds.

### In vivo NP administration and NPD/RPD measurements

All animal procedures were performed in accordance with the guidelines and policies of the Yale Animal Resource Center (YARC) and approved by the Institutional Animal Care and Use Committee (IACUC) of Yale University. Male and female mice homozygous for the F508del mutation (fully backcrossed C57/BL6 background) primarily aged 3-6 months old were used. All mice were genotyped prior to use. Mice were anesthetized using isoflurane. Once respirations reduced to 1 breath/second, NPs were administered IV via retro-orbital injection. 2 mg of NPs were resuspended in 150 μL DPBS and sonicated prior to injection. Four total doses of 2 mg of NPs were given over the course of two weeks. γPNA/DNA NPs contain ~2 μg/0.2 nmol of PNA and ~2 μg/0.1 nmol DNA per mg; each animal received ~0.2 mg/kg of PNA and donor DNA per dose. Nasal potential difference (NPD) and rectal potential difference (RPD) measurements as indicators of ion transport across respiratory and GI epithelia *in vivo* were performed as previously described.^60^ One baseline measurement and up to 3 post-NP-treatment measurements were done for each mouse at least two weeks after the last treatment or measurement. Briefly, mice were anesthetized with ketamine/xylazine, after which an electrode probe is placed into one nostril (NPD) or the rectum (RPD) at 3 mm with a reference electrode with 3% agar in Ringer’s solution placed subcutaneously in the tail. Saline flow through the probing electrode is controlled by a microperfusion pump at 23 μLmin^−1^ for NPD and 0.5 mlhour^−1^ for RPD. Both electrodes are connected to a voltmeter to measure potential differences, which were recorded following a course of solutions: control Ringer’s solution, Ringer’s solution containing 100 μM amiloride, chloride-free solution with amiloride, and then chloride-free solution with amiloride and forskolin/IBMX. The 0 Cl^−^ solution used for all GI tract tissues contained 5 mM barium hydroxide to block potassium currents.

### Ex vivo Ussing measurements, BAL fluid analysis, serum analysis, and histology

Following euthanasia of F508del/F508del mice with a lethal dose of ketamine/xylazine, blood was collected via retro-orbital eye bleed or cardiac puncture, and animals were heart perfused with heparinized DPBS to remove remaining blood from circulation. Aliquots of collected blood were separated into serum and plasma by centrifugation and serum was analyzed for cytokine levels using a bead-based multiplex assay (Milliplex; Millipore) for Luminex. Serum was also collected and sent to Antech Diagnostics for blood chemistry analyses per standard protocols. Lungs were filled with 2 mL DPBS containing protease inhibitors and 0.5M EDTA at pH 8. for bronchoalveolar lavage, and this fluid was collected for further analysis (cell count and cytokine levels). The right lobes of the lung were tied off and the left lobe was collected for histopathology by first inflating with 0.5% low melt agarose at 37 °C at constant pressure and then fixing in 10% neutral buffered formalin solution. Following fixation, lungs were embedded in paraffin, sectioned, and stained with hematoxylin and eosin prior to imaging. Samples of epithelia from nose, ileum, duodenum, jejunum, distal colon, and rectum were mounted in Ussing chambers and Ussing analyses were performed as described above. The 0 Cl^−^ solution used for all GI tract tissues contained 5 mM barium hydroxide to block potassium currents.

### Biodistribution

After IV injection of fluorescent agents, blood collection, and euthanasia as outlined above, animals were perfused transcardially with heparinized DPBS (100 USP/mL). Organs (brain, heart, lungs, liver, spleen, kidneys, bone) were harvested. Fluorescent agent accumulation in the organs was visualized and quantified using an In Vivo Imaging System (IVIS, Perkin Elmer). Tissues were then homogenized into a single-cell suspension through a 70 μm cell strainer and washed twice with PBS by centrifugation and resuspended in PBS containing 2% bovine serum albumin (BSA). Uptake of fluorescent agents in single cells was analyzed by flow cytometry (Attune NxT) and compared to cells harvested from untreated control animals.

### Genomic DNA extraction

Genomic DNA was harvested from cells and tissues using the ReliaPrep gDNA Tissue Miniprep System (Promega) according to the manufacturer’s instructions. Tissues were first homogenized into a single cell suspension prior to gDNA extraction.

### Droplet digital PCR and deep sequencing

Droplet digital PCR (ddPCR) was used to quantify gene editing at the target site. The concentration of genomic DNA (gDNA) samples was measured using the nanodrop and 80-100 ng of gDNA was used for each ddPCR reaction. ddPCR reactions were set up as follows: 11 μL 2×ddPCR™ supermix for probes (no dUTP) (BioRad), 0.2 μL FWD primer (100 μM), 0.2 μL REV primer (100 μM), 0.053 μL CF (HEX) probe (100 μM), 0.053 μL wild-type (edit) probe (100 μM) (Integrated DNA Technologies), 0.5 μL EcoRI, 10 μL gDNA and nuclease-free dH^2^O combined. The primers used were:

FWD: 5’-TGCTCTCAATTTTCTTGGAT-3’
REV: 5’-GGCAAGCTTTGACAACA-3’

The ddPCR probes were:

CF (5’ HEX): 5’-ATCATAGGAAACACCAATGATAT-3’
Wild-type (5’ FAM): 5’-CATCATAGGAAACACCAAAGAT-3’

ddPCR droplets were generated using an automated Droplet Generator (AutoDG™; BioRad). Thermocycler conditions were as follows: 95 °C 10 min, (94 °C 30s, 53.7 °C 2 min – ramp 2 °C/s) x 40 cycles, 98 °C 10 min, hold at 4 °C. Following the PCR reaction, droplets were left at 4 °C for at least 30 minutes and read using the QX200 Droplet Reader (BioRad). ddPCR data was analyzed using QuantaSoft™ software. Data are represented as the fractional abundance of the edited CFTR allele. gDNA from *in vitro* or *in vivo* samples were amplified by PCR to detect both on-target and off-target editing. PCR reactions were performed using KAPA HiFi HotStart ReadyMix (Roche) according to the manufacturer’s instructions. PCR primers for the on-target region were:

FWD: 5’-TCTGCTCTCAATTTTCTTGGA-3’
REV: 5’-GGCAAGCTTTGACAACACTC-3’

Thermocycler conditions were as follows: 95 °C 3 min, (98 °C 20s, 55 °C 15s, 72 °C 15s) x 35 cycles, 72 °C 10 min, hold at 4 °C. To study off-target effects, we looked for alterations in regions of partial homology to both the PNA and the donor DNA. The 11 off-target sites were chosen by searching for sites in the C57BL/6 mouse genome with high (>94%) sequence homology to the PNA binding site (matching 18 bp of the 19 bp PNA binding site). Primers were designed to amplify each of these sites and the PCR amplicons were submitted for deep sequencing (NGS). Primers and barcodes for off-targets are shown in **Supplementary Table 2**. PCR products were purified using the QIAquick PCR Purification Kit (Qiagen). PCR products were prepared for NGS by end-repair and adapter ligation according to Illumina protocols and samples were sequenced by either the the Illumina NovaSeq 6000 with 100 or 150 paired-end reads at the Yale Center for Genome Analysis (YCGA). This instrument has a Q30 score of >85%, indicating that at least 85% of the bases have an error rate of 0.1% or less. This value is an average across the whole read length, and error rate increases towards the end of the reads. Sequencing data was mapped to corresponding reference sequences and analyzed using CRISPResso2. Potential PCR artifacts of the donor DNA were eliminated by using AMPure XP beads (Beckman Coulter) to purify genomic DNA according to the manufacturer’s instructions prior to PCR amplification in a subset of samples.

### Data analysis

Results were analyzed using GraphPad Prism (version 8.1.2). Data are presented as individual data points or as mean ± standard error of the mean (s.e.m.). To compare *in vitro* Ussing readouts and ddPCR editing frequencies in ALI cultures, we used ANOVA with Dunnett’s post-test for multiple comparisons. To compare *in vitro* and *in vivo* off-target effects analyzed by NGS and *in vivo* ddPCR editing frequencies among treatment conditions, we used the Wilcoxon rank sum test on pooled data across off-target sites and multiple organs, respectively. We used the Mann-Whitney test to compare NPD, RPD, Ussing and BAL readouts following *in vivo* treatment among treatment conditions. To compare pre- and post-treatment *in vivo* NPD and RPD values, we used the Wilcoxon matched-pairs signed rank test. For all statistical tests, p<0.05 was considered to be statistically significant.

## ABBREVIATIONS

ALI: air-liquid interface
BAL: bronchoalveolar lavage
CF: cystic fibrosis
CFTR: cystic fibrosis transmembrane conductance regulator
ddPCR: droplet digital PCR
DCM: dichloromethane
DLS: dynamic light scattering
GI: gastrointestinal
HR: homology-dependent repair
IV: intravenous
IVIS: In Vivo Imaging System
NECs: nasal epithelial cells
NER: nucleotide excision repair
NGS: next generation sequencing
NP: nanoparticle
NPD: nasal potential difference
PLGA: poly(lactic-co-glycolic acid)
PNA: peptite nucleic acid
RPD: rectal potential difference
scRNA-seq: single-cell RNA sequencing
SEM: scanning electron microscopy
TFO: triplex-forming oligonucleotide
UMAP: Uniform Manifold Approximation and Projection

## Supplementary Information

**Supplementary Table 1.** Characterization data for PLGA NP formulations.

| Formulation | Diameter (nm) | Polydispersity Index (PDI) | Zeta Potential (mV) |
| --- | --- | --- | --- |
| PLGA-Cy5 | $257 \pm 3$ | 0.050 | $-10.7 \pm 1.4$ |
| PLGA DiD | $273 \pm 2$ | 0.190 | $-17.2 \pm 0.7$ |
| PLGA Blank | $240 \pm 3$ | 0.110 | $-23.4 \pm 0.5$ |
| PLGA<br>PNA/DNA | $265 \pm 6$ | 0.110 | $-27.6 \pm 1.0$ |
| PLGA<br>γPNA/DNA | $265 \pm 3$ | 0.130 | $-28.2 \pm 0.3$ |

**Supplementary Table 2.**
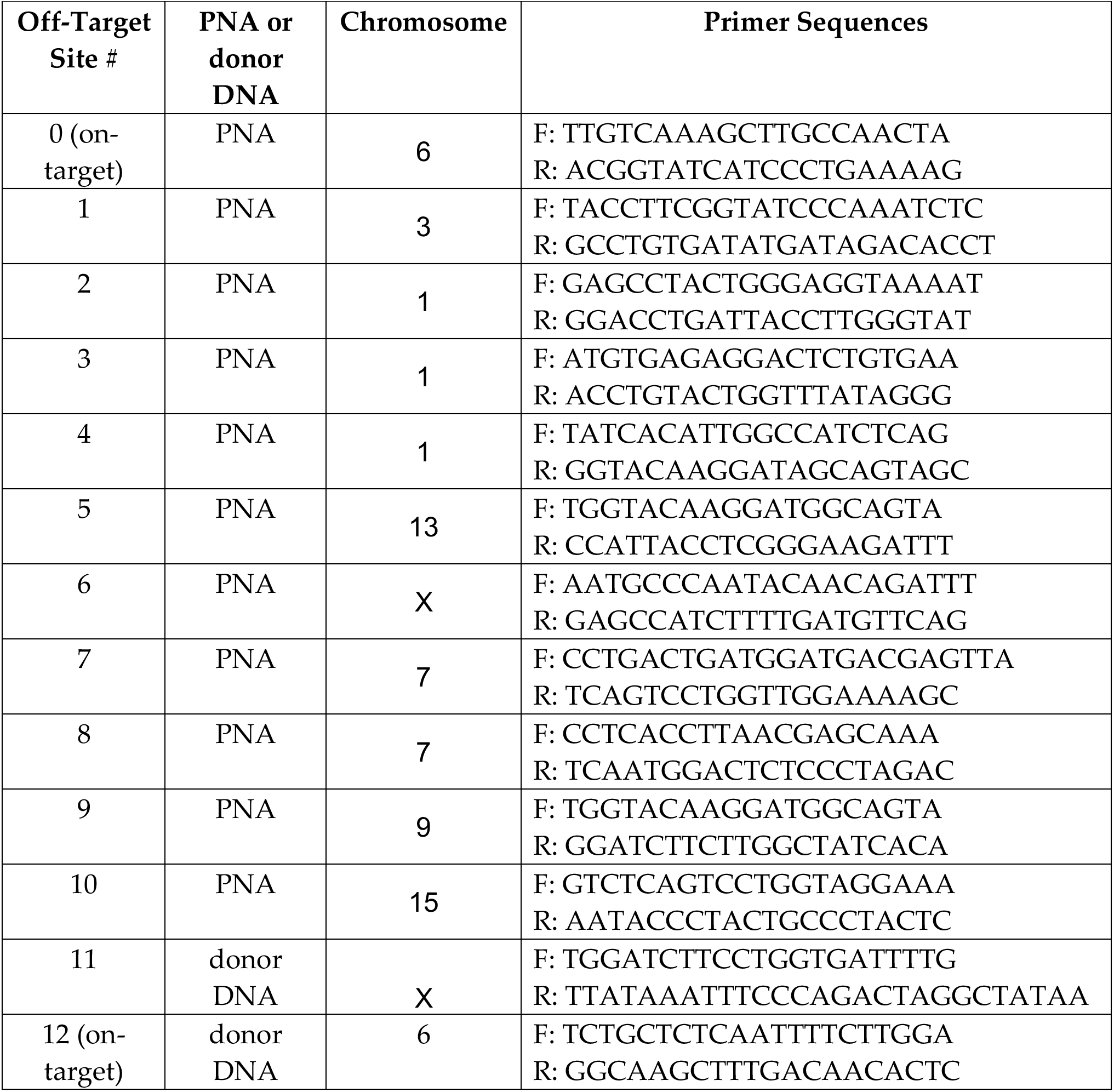
Primer Sequences for PNA and donor DNA off-target analysis.

**Supplementary Table 3.** Blinded histopathology analysis of key organs following blank or γPNA NP treatment in C57BL/6J mice. 1 = no lymphoid aggregates, 2 = at least 1 lymphoid aggregate. N=3 animals per group.

| **Treatment Group** | **Average Score** |  |  |  |
| --- | --- | --- | --- | --- |
|  | **Lungs** | **Liver** | **Spleen** | **Kidneys** |
| Untreated Control | 1.33 | 2 | 1 | 1 |
| Blank NPs | 1 | 1.33 | 1 | 1 |
| γPNA/DNA NPs | 1.67 | 1 | 1 | 1 |

**Figure S1.**
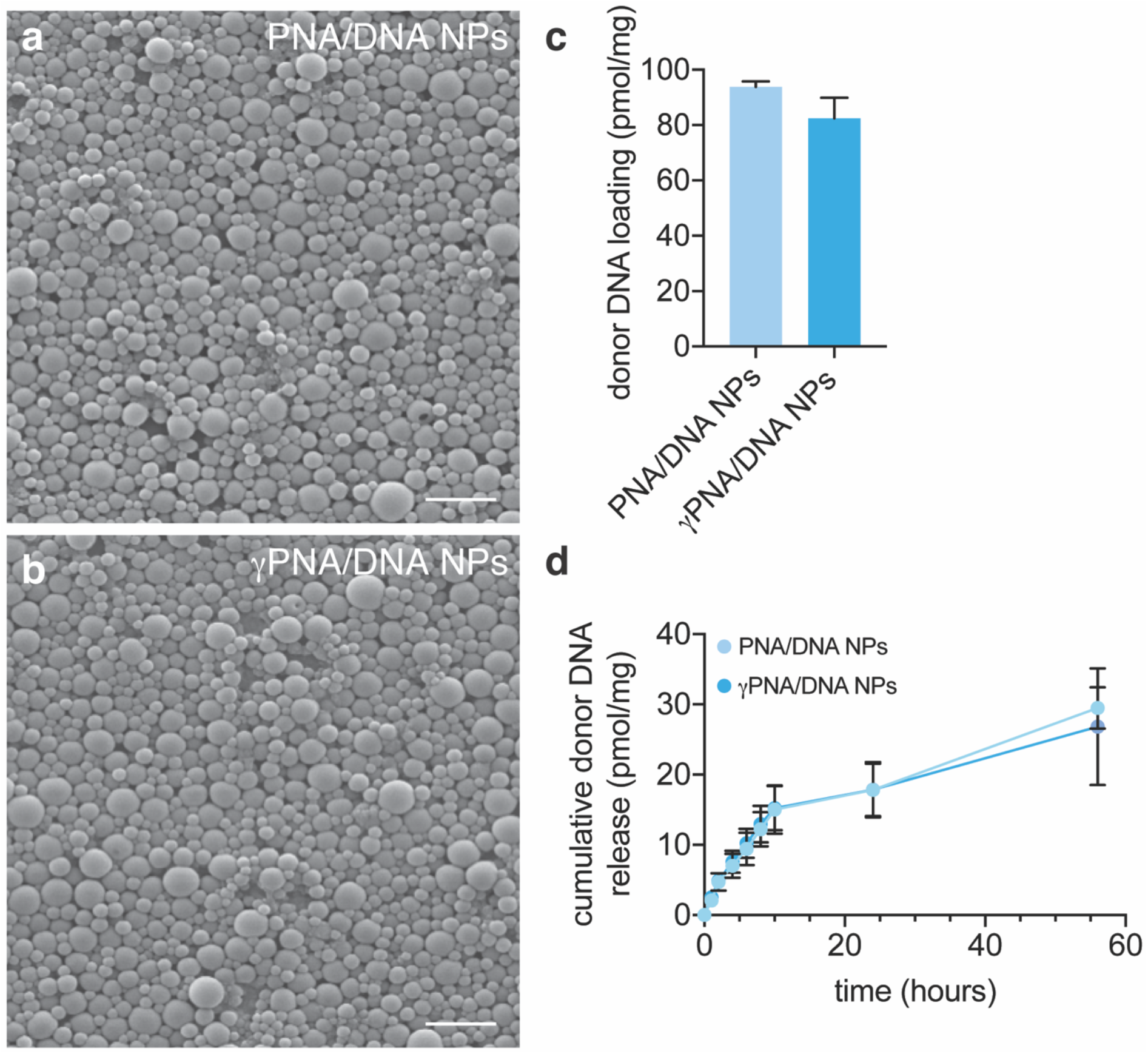
Characterization of NP formulations. SEM images of PLGA NPs containing (a) PNA and donor DNA and (b) γPNA and donor DNA. Scale bars, 1 μm. (c) Total donor DNA loading of PNA/DNA and γPNA/DNA PLGA NPs as measured by the Oligreen assay. (d) Cumulative donor DNA release from PNA/DNA and γPNA/DNA PLGA NPs as measured by the Oligreen assay.

**Figure S2.**
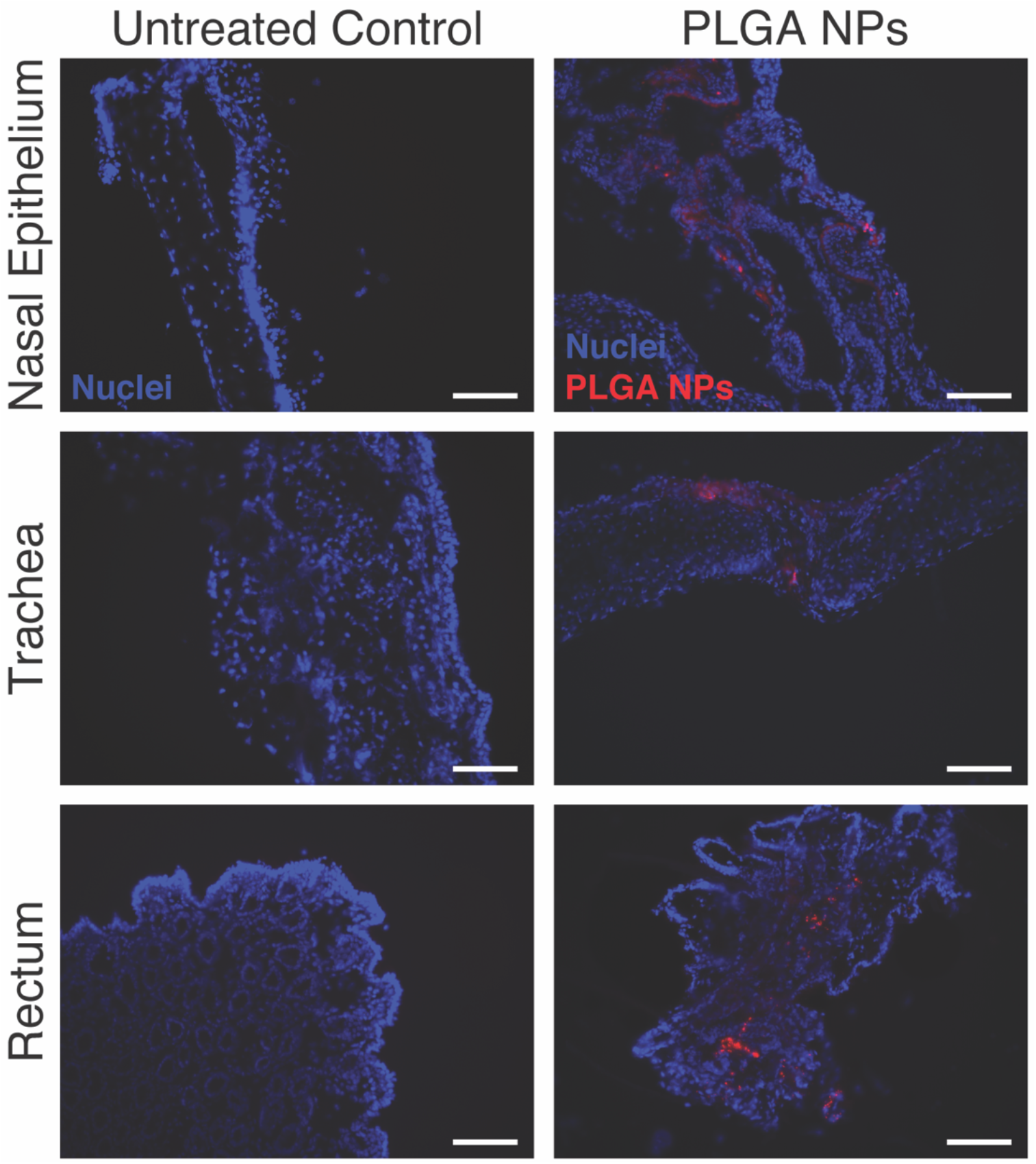
Biodistribution of PLGA NPs to the nasal epithelium, trachea, and rectum. Representative fluorescence microscopy images indicating biodistribution of DiD-loaded PLGA NPs (red) to the nasal epithelium, trachea, and rectum, at 24 hours post-IV administration *in vivo*, with untreated control images provided for comparison. Cell nuclei are shown in blue. Scale bars, 100 μm.

**Figure S3.**
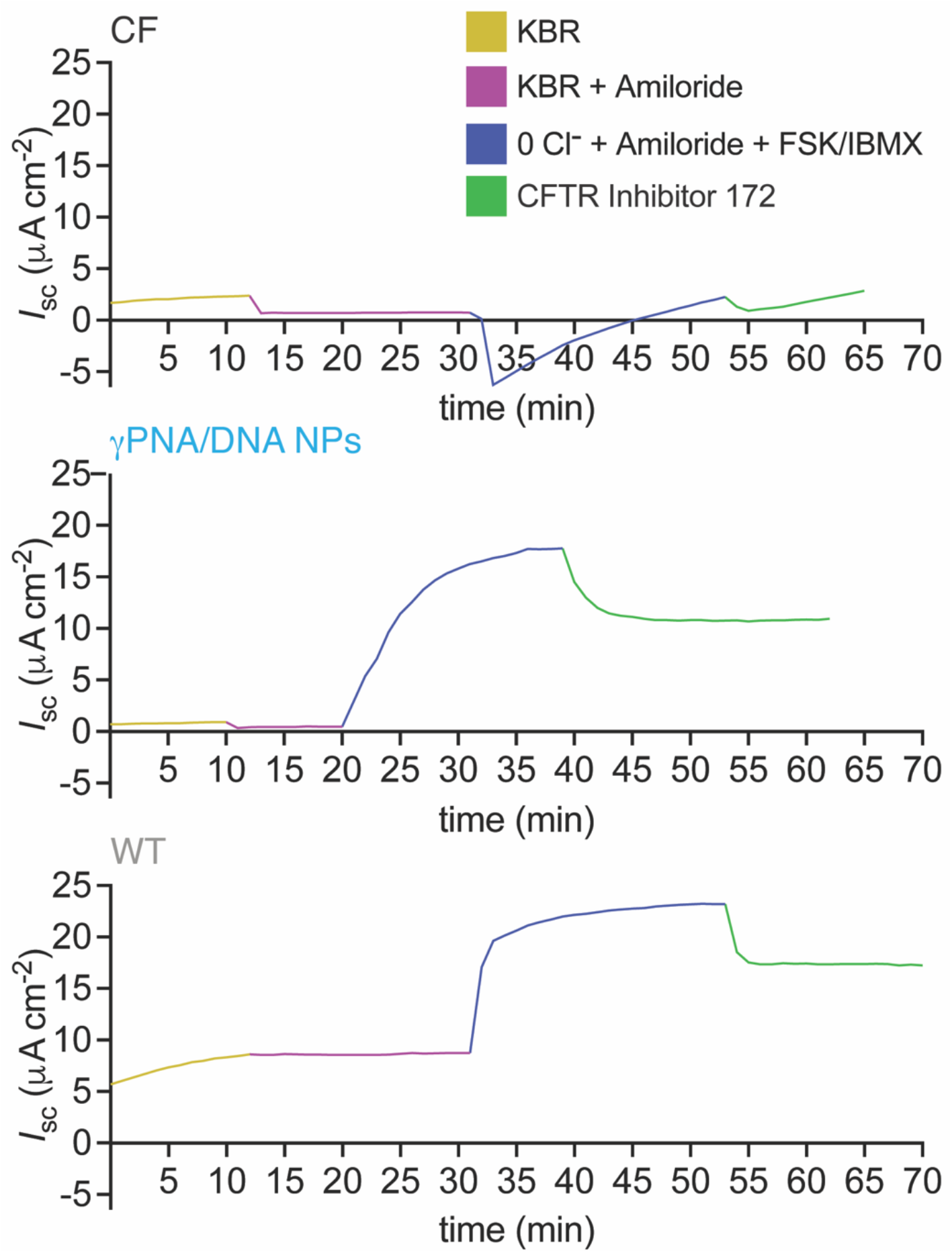
Representative NEC ALI Ussing Traces. Primary NEC ALI cultures were loaded into Ussing chambers and short-circuit current (*I_sc_*) was recorded during the addition of amiloride, forskolin/IBMX, and a CFTR inhibitor (172). Representative traces of CF NEC ALI cultures, CF NEC ALI cultures treated with γPNA/DNA NPs, and wild-type NEC ALI cultures are shown.

**Figure S4.**
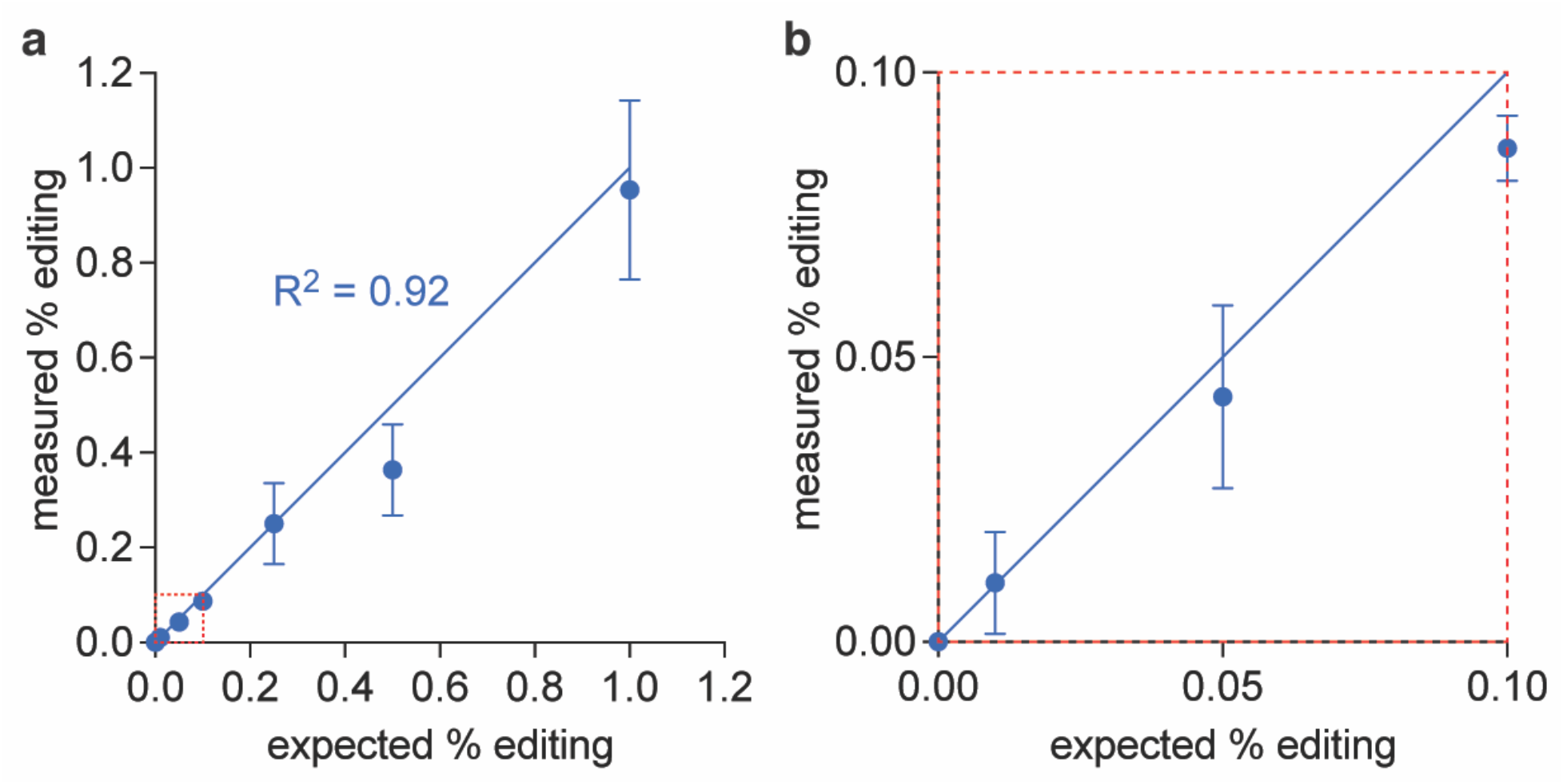
CF F508del Droplet digital PCR (ddPCR) assay validation. A ddPCR assay was designed in which fluorescent probes differentiating two alleles (wild-type or mutant (F508del) are specific for the gDNA template present in the PCR reaction. (a) The expected percent fractional abundance of the wild-type allele (wild-type/wild-type + mutant) is plotted against the measured fractional abundance for an experiment in which increasing known amounts (0.01% to 1% by weight) of wild-type gDNA were spiked into samples of mutant F508del gDNA. The observed correlation is linear with an R^2^ value of 0.92. (b) An expanded view of the lower end of the analysis range indicated by the red dashed box in panel (a). Error bars indicate SD.

**Figure S5.**
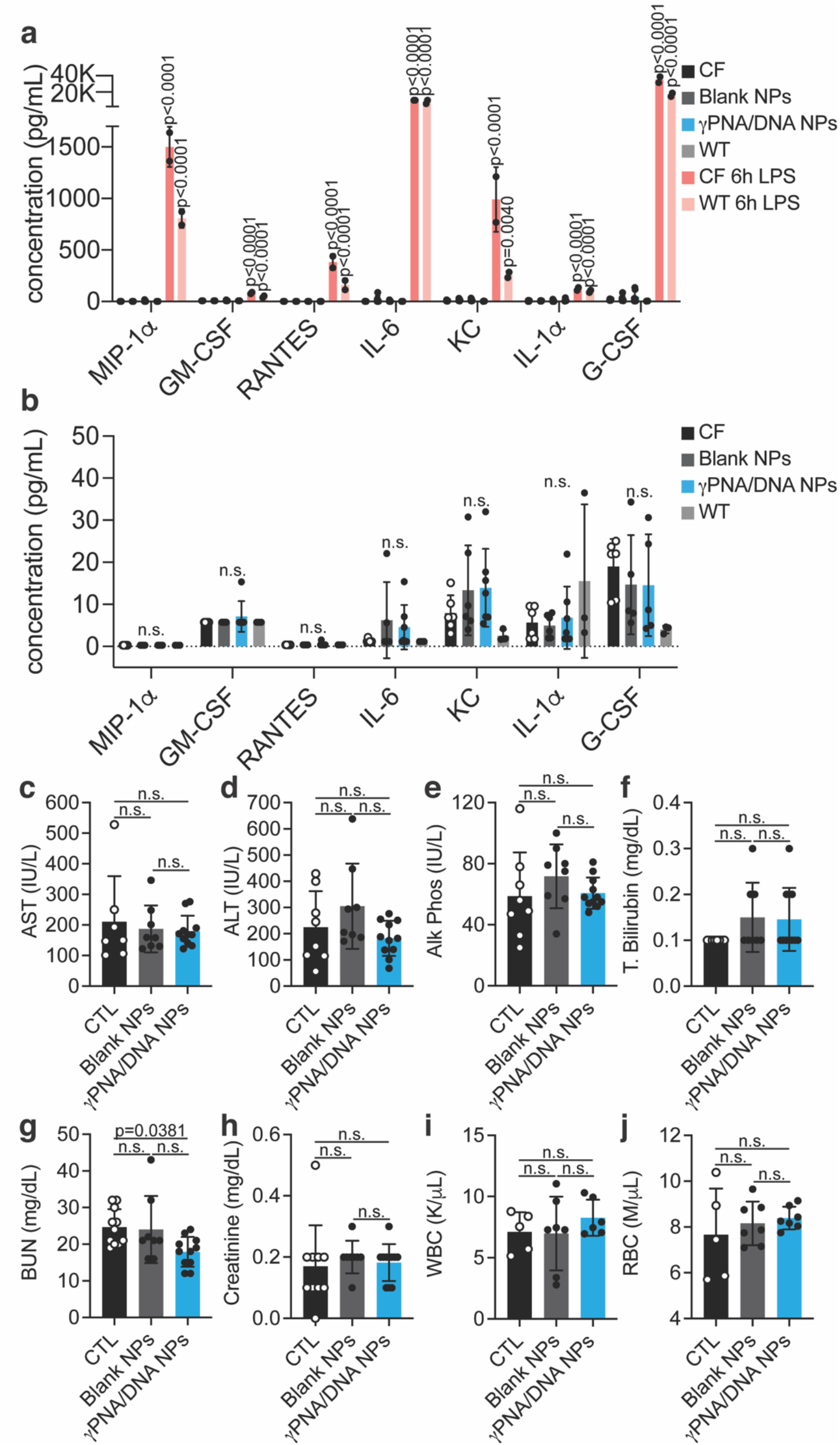
PNA NPs are non-toxic and do not elicit an immune response. (a) Cytokine production in BAL fluid of CF (untreated control), blank NP-treated mice, γPNA NP-treated mice, CFTR KO mice nebulized with LPS (positive control), and WT mice nebulized with LPS (positive control). (b) Same plot as in (a) without positive LPS controls. (d)-(k) Blood serum chemistry analyses for untreated control, blank NP-treated, and γPNA NP-treated mice.

**Figure S6.**
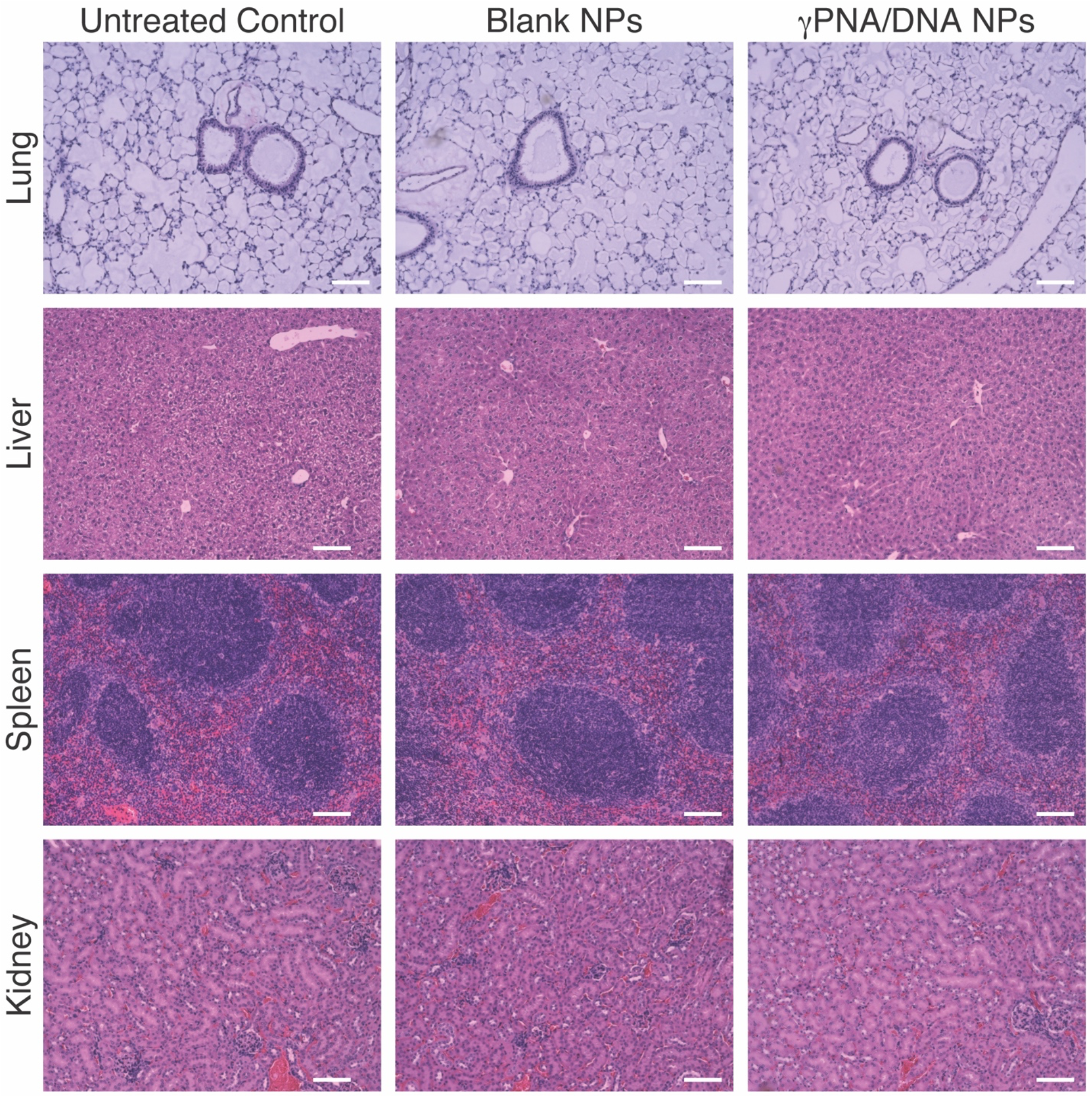
Blank and γPNA/DNA NP-treated mice exhibit normal histology in the lung, liver, spleen, and kidneys. Histology images of untreated, Blank NP-treated, and γPNA/DNA NP-treated mouse lungs, livers, spleens, and kidneys that were paraffin-embedded and stained with haematoxylin and eosin. Scale bars, 100 μm.

**Figure S7.**
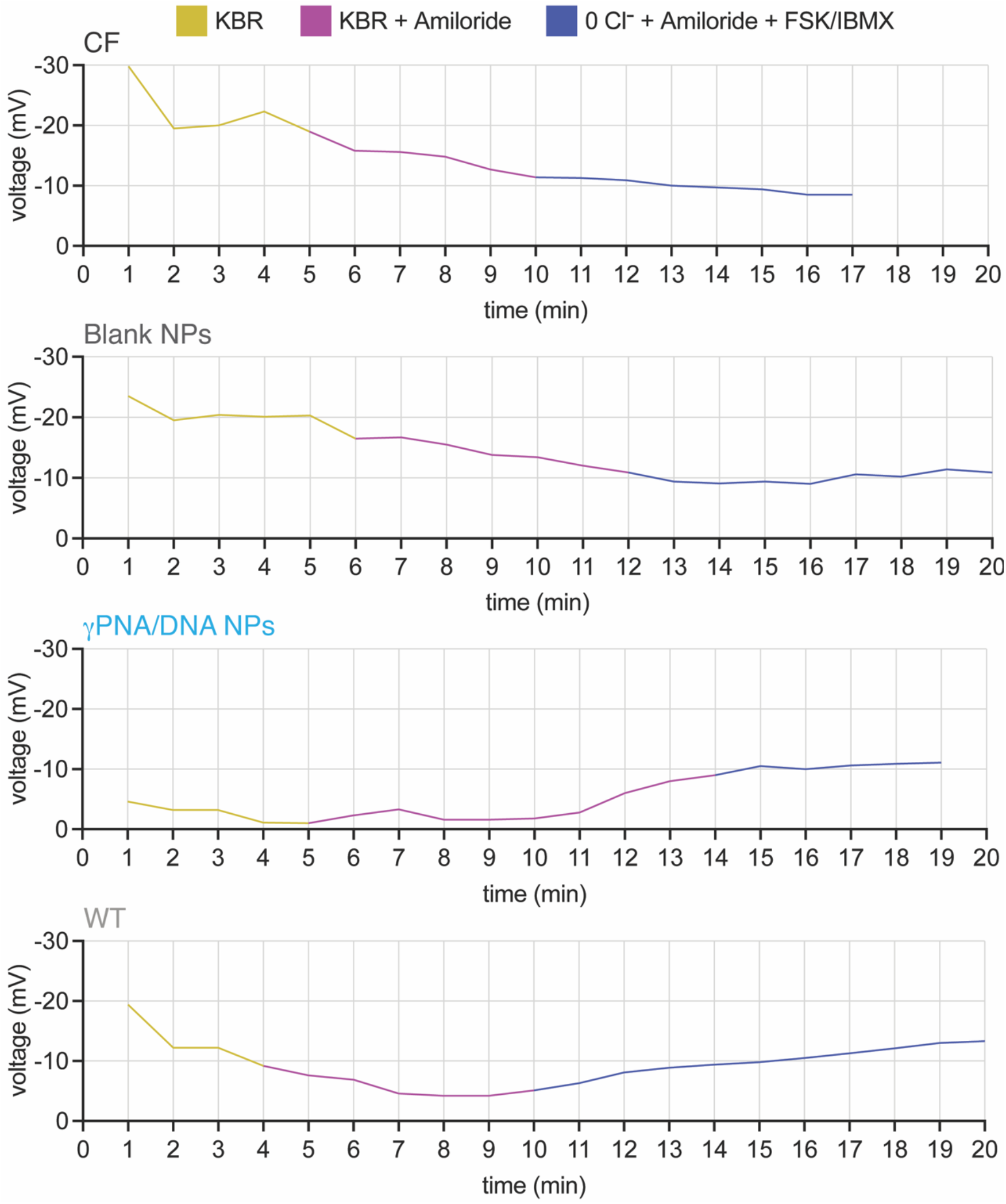
Representative Raw NPD Traces. Nasal epithelia of F508del-CFTR mice were probed with an electrodes connected to a voltmeter to measure potential differences recorded following a course of solutions: Ringer’s (KBR), KBR with amiloride, and chloride-free solution with amiloride and forskolin/IBMX. Representative traces of untreated CF mice, CF mice treated with Blank NPs or γPNA/DNA NPs, and wild-type mice shown.

**Figure S8.**
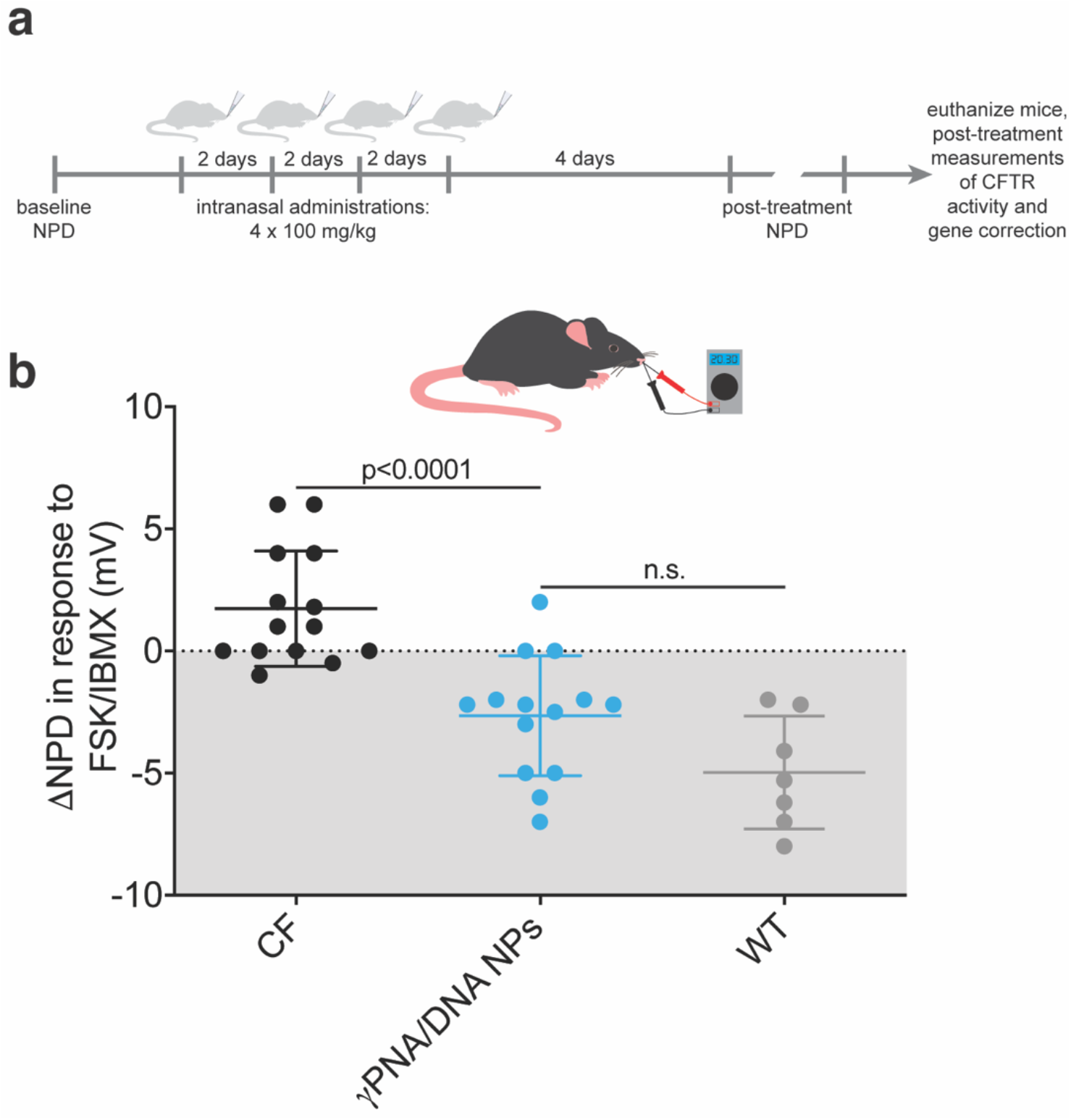
Local intranasal γPNA NP treatment results in robust *in vivo* NPD response. (a) Schematic of *in vivo* intranasal dosing scheme for PLGA/PBAE/MPG NPs loaded with γPNA and donor DNA. (b) NPD measurements following γPNA/DNA NP treatment with CF (black circles) and wildtype (light grey circles) controls. Grey region indicates wild-type range.

## Acknowledgements

We thank biostatisticians Dr. Daniel Zelterman and Dr. Wei Wei for their assistance with statistical analyses and Trucode Gene Repair Inc. for providing a portion of the PNAs used in this study. This work was supported by grants from the National Institutes of Health (NIH; UG3 HL147352, R01 HL125892) and the Cystic Fibrosis Foundation (CFF; EGAN641558). A.S.P. was supported by two NIH NRSAs (a T32 GM86287 training grant and an F32 HL142144 individual postdoctoral fellowship), a postdoctoral research fellowship award from the CFF (PIOTRO20F0), a K99/R00 Pathway to Independence award from the NIH (K99 HL151806), and a Postdoc-to-Faculty Transition Award from the CFF (PIOTRO21F5).

## Author Contributions

A.S.P., C.B., P.M.G., W.M.S., and M.E.E. designed the experiments. A.S.P., P.M.G., W.M.S., and M.E.E. wrote the manuscript. A.S.P., C.B., C.L., Y.D., and D.W. performed the experiments. A.S.P., Y.D., and M.E.E. analyzed the data and prepared the figures. All authors discussed the results and assisted during manuscript preparation.

